# Repurposing PeTriBERT for Protein–Protein Interaction Prediction with Sequence–Structure Fusion

**DOI:** 10.64898/2026.09.29.755338

**Authors:** Arthur Wavreille, Gabriel Krouk, Baldwin Dumortier

## Abstract

Recent advances in artificial intelligence have enabled models to capture protein sequence and structural features. We present PeTriPPI2, a hybrid framework for protein–protein interaction (PPI) prediction that combines sequence embeddings from ESM-2 with structural representations from PeTriBERT, an encoder originally designed for inverse folding. We evaluate this sequence–structure approach on the Pinder dataset. On a test set of 2,342 protein pairs, the selected checkpoint achieves 0.898 accuracy, 0.917 precision, 0.876 recall, and 0.896 F1 score for the interacting class (AUROC 0.963). Compared with SpatialPPIv2 on the same test data, PeTriPPI2 has higher precision (0.917 versus 0.886) but lower recall (0.876 versus 0.942) and F1 score (0.896 versus 0.913). Component ablations indicate that both the sequence and structural pathways contribute to this configuration.

## 1 Introduction

Proteins are fundamental macromolecules that sustain virtually all cellular processes, from metabolic regulation and immune response to signal transduction and structural organization ([1]). Their functions are dictated by their amino acid sequences and the resulting three-dimensional conformations, which govern their chemical properties, binding specificity, and dynamic behavior ([2]). Understanding protein function has therefore long been a central challenge in molecular biology, with profound implications for medicine, agriculture, and biotechnology.

A critical aspect of protein function lies in their ability to physically interact with one another. Protein–protein interactions (PPIs) form the backbone of molecular networks, enabling enzymatic complexes, signal cascades, and structural assemblies. Characterizing PPIs is essential for elucidating biological mechanisms, annotating uncharacterized proteins, and identifying therapeutic targets. However, despite the development of high-throughput experimental techniques such as yeast two-hybrid screens, affinity purification, and cross-linking mass spectrometry, large-scale mapping of PPIs remains limited by cost, labor intensity, and incomplete coverage ([3]). This gap has motivated the rise of computational methods aimed at predicting PPIs accurately and at scale.

Over the past decade, artificial intelligence (AI) has dramatically advanced protein science, particularly through breakthroughs in protein structure prediction. The release of AlphaFold2 ([4]) and RoseTTAFold ([5]) marked a turning point by solving the protein folding problem with unprecedented accuracy, often rivaling experimental methods. These deep-learning architectures combined evolutionary information through multiple sequence alignments (MSAs) with geometric reasoning and transformer attention mechanisms, providing atomic-level models of protein structures. Their success inspired a wave of extensions beyond monomer folding, including AlphaFold-Multimer ([6]) and algorithms such as FoldDock ([7]) or Alphafold Initial Guess [8], which adapted AlphaFold2 outputs to score and predict protein–protein complexes. In particular, Alphafold Initial Guess is now widely adopted and used ([9, 10, 11, 12, 13, 14, 15, 16]). Parallel efforts, such as ESMFold ([17]) or ProtTrans [18], have demonstrated that large protein language models trained on billions of sequences can predict structures directly from single sequences, further broadening the accessibility and scalability of structure prediction. Collectively, these advances have reshaped structural biology and created new opportunities to tackle downstream challenges such as interaction prediction, docking, and protein design.

In parallel to structural modeling, the last few years have witnessed rapid progress in sequence-based learning which progressed quikly by the mean of geometric/equivariant architecture [19, 20, 21, 22]. Protein language models, inspired by natural language processing, leverage the vast availability of amino acid sequences to learn contextual embeddings that capture structural and functional properties. Early deep learning methods such as DeepPPI ([3]) used neural networks with hand-crafted sequence descriptors to classify interacting pairs, while more sophisticated architectures like D-Script ([23]) and Topsy-Turvy ([24]) introduced transfer learning and pre-trained embeddings to improve performance. The introduction of transformer-based language models such as ProtBert, ProtT5 ([18]), and ESM-2 ([25]) set new standards by encoding evolutionary and structural information directly from raw sequences. These models have achieved remarkable success across multiple protein-related tasks, including annotation, mutational effect prediction, and PPI detection. Nevertheless, sequence-only approaches remain limited by their inability to explicitly model the three-dimensional compatibility of protein surfaces, which is central to physical binding.

To address these limitations, structure-based methods have been developed, leveraging experimentally determined or predicted structures to model protein interactions at the geometric level. Approaches such as Struct2Graph ([26]), GraphPPIS ([27]), and DeepInteract ([28]) represent proteins as graphs or point clouds, and apply graph neural networks or geometric transformers to capture spatial organization and interface-specific features. More recent developments, such as PeSTo ([29]), directly process atomic coordinates using equivariant architectures, ensuring that predictions remain consistent under rigid-body transformations. While structure-based approaches provide interpretability and higher specificity in modeling binding interfaces, their adoption was long constrained by the scarcity of structural data. The wide availability of high-quality predicted structures following AlphaFold2 has significantly alleviated this bottleneck, enabling the scaling of structure-based PPI prediction.

Recognizing the complementary strengths of sequence and structural representations, several recent studies have explored hybrid approaches that combine both modalities. These methods generally leverage protein language models for scalable sequence embeddings while incorporating structural encoders to capture geometric features relevant to binding. For instance, SpatialPPIv2 ([30]) follows this direction by integrating sequence embeddings with structural graph representations for PPI prediction.

In this work, we pursue a similar hybrid strategy but with a distinct design that leverages pre-trained models with demonstrated versatility. We introduce PeTriPPI2, which combines the ESM protein language model for sequence representation with PeTriBERT, a transformer-based structural encoder originally developed for the inverse folding problem ([31]). We evaluate whether a pretrained structural encoder can contribute to interaction prediction when combined with ESM-2. On the Pinder benchmark dataset ([32]), the combined model reaches 0.896 F1 for the interacting class, with a higher precision and a lower recall than SpatialPPIv2 on the same test data. Ablations provide evidence that both pathways contribute to this model configuration.

## 2 Method

PeTriPPI2 is a hybrid sequence–structure model, leveraging the representational power of two pretrained encoders: ESM-2, a widely used protein language model, and PeTriBERT (specifically its PeTriMPOV variant), a protein sequence–structure encoder originally designed for inverse folding. Figure 1 summarizes the PeTriPPI2 architecture. For each protein pair, the input structures are used to construct residue-level structural features and a pairwise attention bias. The two amino acid sequences receive token and segment embeddings before passing through a five-layer transformer encoder whose attention incorporates this structural bias. In parallel, a frozen ESM-2 model produces sequence representations. The transformer outputs and dimension-adapted ESM-2 representations are concatenated and mean-pooled; a feed-forward prediction head followed by a sigmoid produces an interaction probability. The model is fine-tuned on the Pinder dataset while keeping the ESM-2 weights frozen.

**Figure 1:**
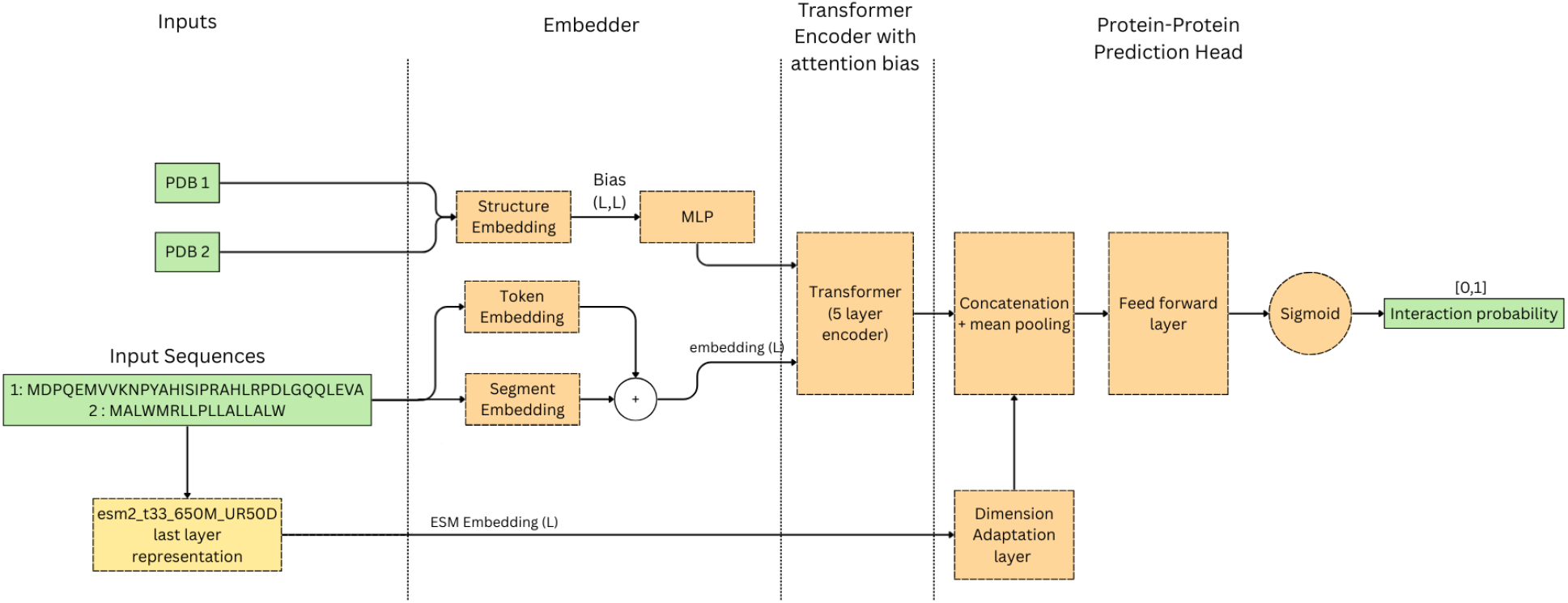
Overview of PeTriPPI2. Structural information from the two input proteins supplies the attention bias of the PeTriBERT/PeTriMPOV encoder, while their sequences supply token and segment embeddings. Frozen ESM-2 representations form a parallel sequence pathway. The two pathways are combined before a prediction head outputs the probability of interaction.

### 2.1. Dataset

High-quality and reliably labeled protein–protein interaction data are critical yet difficult to obtain for PPI prediction. Assessing biological interactions varies considerably with the chosen experimental method (*in vivo* or *in vitro*) and is subject to experimental uncertainty (e.g., protein synthesis quality, temperature, contamination, etc.). Moreover, datasets typically contain few negative (non-interacting) pairs, as most entries correspond to observed interactions.

This limitation affected our previous approach [33], which relied on the Propedia dataset (20,000 experimentally validated interactions) and required augmentation to generate negative pairs. Since protein–protein interactions are a sparse phenomenon ([34, 35, 36]), unobserved pairs are generally assumed to be negative. While often valid, this assumption does not eliminate the risk of mislabeling.

For the present work, we used the PINDER dataset, a larger and more rigorously curated resource for protein interactions. PINDER integrates structural data from both the RCSB Protein Data Bank (PDB) and the AlphaFold Database, ensuring broad coverage and diversity. To avoid data leakage, the dataset was organized through interface-based clustering: structurally similar proteins were grouped into the same cluster and consistently assigned to the same split (train/validation/test). This strategy provides more reliable validation and test results.

The version of PINDER employed here comprises several million protein dimers, making it particularly suitable for training transformer-based deep learning models, which benefit from large-scale datasets. We additionally excluded PDB files containing only C*α* atoms, since our model requires N, C*α*, and C*β* atoms. After filtering, the training split contained 1,603,337 protein pairs, including 801,553 interacting pairs and 801,784 non-interacting pairs, thereby preserving a balanced 50/50 distribution of positive and negative labels.

### 2.2. Sequence feature extraction with protein language models

ESM-2 was used as a frozen feature extractor. Pretrained language models have been applied to a variety of protein-related tasks [25, 18]. In our framework, ESM-2 complements PeTriBERT, which predominantly captures structural rather than sequential order information [31].

Specifically, PeTriPPI2 employs the final hidden representations from ESM-2. We used the _t33_650M_UR50D model, which contains 33 transformer layers, 650 million parameters, and an embedding dimension of 1280. Since the embedding dimensions of ESM-2 and PeTriBERT differ, we applied a fully connected projection layer to downsample the ESM output to the dimensionality of the PeTriBERT embeddings.

### 2.3. Sequence–structure encoding with PeTriBERT

PeTriBERT [31], and its extension PeTriMPOV [33], is a five-layer, twelve-head transformer encoder trained with masked language modeling (MLM) on protein sequences. Residues are tokenized individually, with additional tokens for sequence boundaries (start, end, pad, and mask).

Unlike conventional language models, PeTriBERT integrates structural information by encoding residue frames (as defined in [4]). These frames are affine orthonormal bases anchored at the backbone atoms of each residue *i* (N, C*α*, and C), computed using a Gram–Schmidt procedure. Mathematically, frames are defined by a rigid transformation *T*_*i*_ that defines the operation used to align the global orthonormal basis with the local residue basis.

In PeTriMPOV, structural frames are integrated into the attention mechanism through three steps:

1. computing relative frames *T*_*i,j*_ (distance vectors and rotation matrices) between all residue pairs *i, j*;

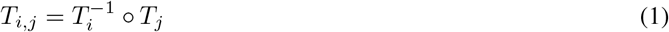

where 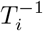 designates the inverse rigid transformation in the mathematical group of rigid transformations, and designates the composition operation in the same group
2. Transforming the relative frames with a fully connected network to produce an (*L, L, h*) tensor (where *L* is the sequence length and *h* the number of attention heads). Each frame *T*_*i*_ are parametrised by 6 parameters, centroid and angles where angles are the coefficients of the rotation matrix are compressed into Euler angles. These parameters are processed by a small fully connected layer to compute a bias matrix **B**.
3. Using this tensor as an attention bias in the transformer. In the transformer, this matrix bias is then added to the attention scores in the attention mechanism:

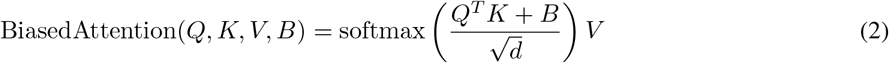

This approach is a rephrasing of the results from ([19]). This transformation acts as an induction bias as it induces a structural representation that is invariant to global rigid transformation which improves generalization throught fighting the curse of dimensionality. This can is easily proved by the following, assuming any global rigid transformation *T*_global_ is applied to the structural data data :

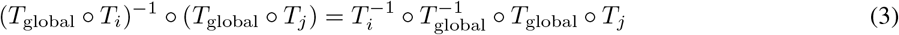

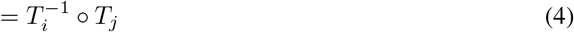

Practically, this transformation is expressed using the centroid **c**_*i,j*_ and rotation matrix *R*_*i,j*_:

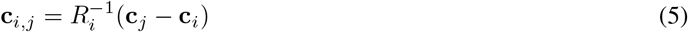

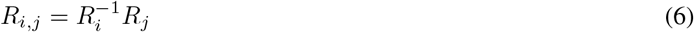

This featurization and attention bias is illustrated on figure 2.

**Figure 2:**
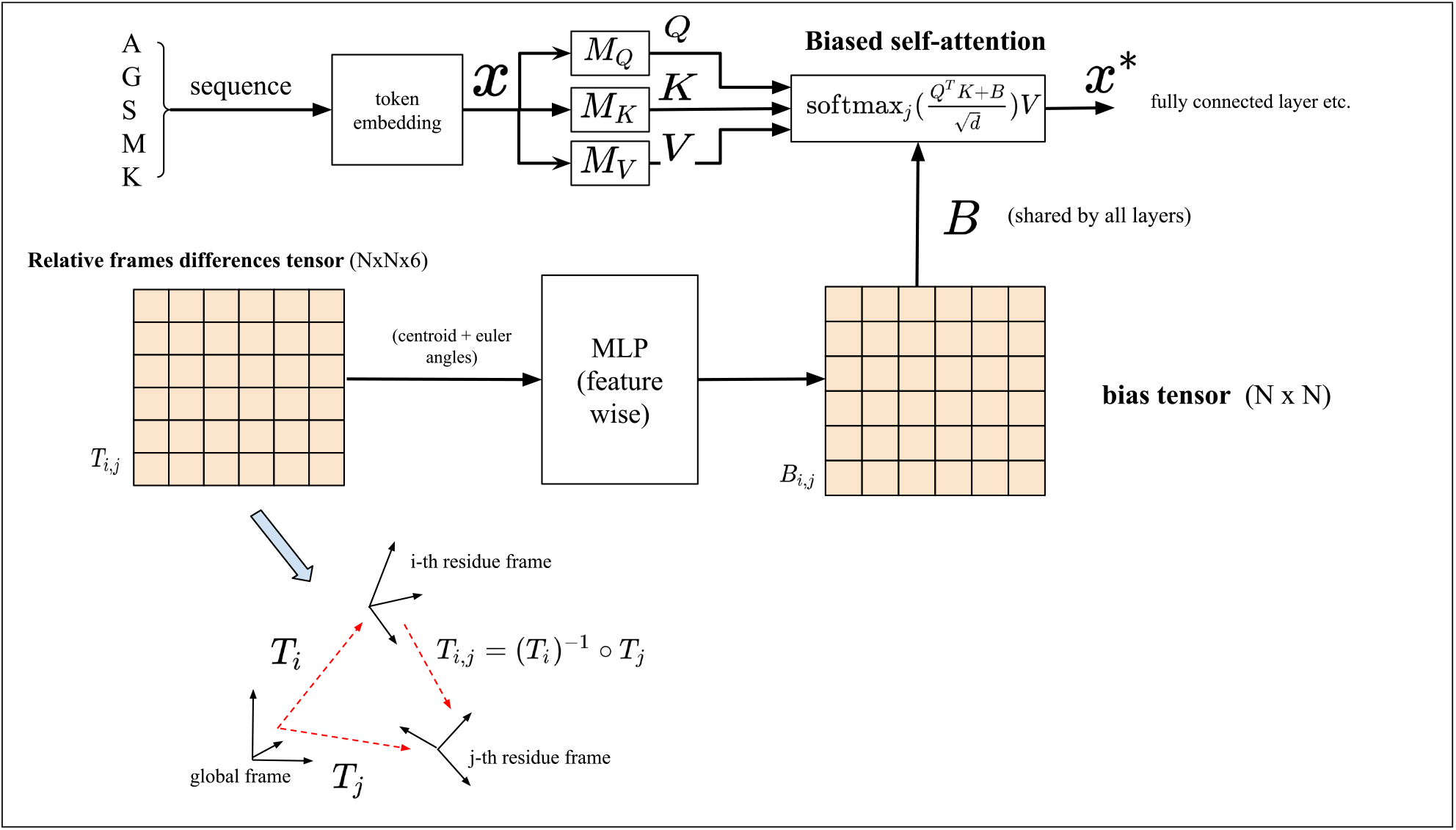
General architecture of PeTriPPIv2.

It parallels the row-wise gated self-attention with pair bias module from AlphaFold2’s Evoformer, though with notable differences: PeTriMPOV operates on a single sequence with its associated structure, rather than on multiple sequence alignments or template-based features.

An additional distinction is that PeTriMPOV does not encode linear sequence position. As shown in our earlier work [31], spatial positioning of residues is more informative than linear indices, and positional encodings do not improve performance.

PeTriMPOV was pretrained on approximately 10^6^ proteins from the AlphaFold Database [37].

### 2.4. Model adaptations

Since PeTriBERT/PeTriMPOV was pretrained on monomeric proteins, several adaptations were required to extend it to protein pairs. First, a segment embedding module was introduced to distinguish between the two input proteins and bias the attention mechanism accordingly. Second, to predict interactions without prior knowledge of quaternary structure (i.e., without predefined relative orientations between the proteins), we removed the anti-diagonal elements of the relative-frame attention bias from gradient propagation. The treatment of inter-chain relative-frame features requires further specification.

Finally, the outputs of PeTriBERT and ESM-2 were concatenated, mean-pooled across the sequence dimension, and passed through a classification head (linear layer followed by sigmoid activation).

## 3 Results

To limit overfitting, we finetuned our model using small batch size and learning rate. We used a simple hyperparameter grid-search. The best run uses, the AdamW optimizer using mini-batch of 32 pairs, and performing validation steps every 1000 batches, using a learning rate schedule with 1000 steps warmup stage and then a linear decay up to 5000 iterations, from which the learning rates remains constant until the end of the training phase.

### 3.1. Comparison with published methods

Table 1 compares PeTriPPI2 with published PPI prediction models, including SpatialPPIv2, FoldDock, Topsy-Turvy, D-script, Struct2Graph, and GNN-PPI, as reported from the SpatialPPI paper [30]. The reported metrics include accuracy, precision, recall, and F1 score. According to the authors, all methods were evaluated on the same test data. For PeTriPPI2, the reported values are recalculated from the confusion matrix of the selected checkpoint (1,030 true positives, 146 false negatives, 1,073 true negatives, and 93 false positives; 2,342 test pairs). Precision, recall, and F1 refer to the interacting class.

**Table 1:** Reported scores for PeTriPPI2 and comparator methods on the same test data, as confirmed by the authors. PeTriPPI2 values are calculated from the selected checkpoint confusion matrix; comparator values are reported in [30]. Precision, recall, and F1 are for the interacting class.

|  | **PeTriPPI2** | SpatialPPIv2 | FoldDock | Struct2Graph | GNN-PPI | Topsy-Turvy | D-script |
| --- | --- | --- | --- | --- | --- | --- | --- |
| Accuracy | 0.898 | **0.910** | 0.888 | 0.812 | 0.771 | 0.774 | 0.704 |
| Precision | **0.917** | 0.886 | 0.899 | 0.811 | 0.730 | 0.839 | 0.857 |
| Recall | 0.876 | **0.942** | 0.875 | 0.813 | 0.858 | 0.678 | 0.490 |
| F1 Score | 0.896 | 0.913 | 0.887 | 0.812 | 0.789 | 0.750 | 0.624 |

### 3.2. Component ablations

The selected checkpoint has an AUROC of 0.963 and an AUPRC of 0.946. In the supplied single-component ablations on the same 2,342 test pairs, removing ESM-2 yields 0.816 accuracy and 0.820 F1 for the interacting class (AUROC 0.900), while removing the PeTriBERT pathway yields 0.523 accuracy and 0.118 F1 (AUROC 0.505). These comparisons indicate that both pathways contribute to this configuration, although the ablation checkpoints alone do not establish statistical variability across training runs.

## 4 Discussion and Conclusion

We introduced PeTriPPI2, a hybrid framework for protein–protein interaction prediction that fuses sequence features from a large protein language model (ESM-2) with structure-aware representations from a pretrained encoder (PeTriBERT/PeTriMPOV). The selected checkpoint yields 0.898 accuracy and 0.896 F1 for the interacting class on 2,342 test pairs; its AUROC is 0.963. The ablation results are consistent with a contribution from both pretrained pathways to this configuration. This supports investigating structural encoder transfer beyond the original inverse-folding task.

Compared with the SpatialPPIv2 scores in Table 1, PeTriPPI2 exhibits higher precision (0.917 versus 0.886), lower recall (0.876 versus 0.942), and lower F1 (0.896 versus 0.913). This operating point may be useful when false positives are costly, but the comparison does not establish an advantage across all application settings. Threshold selection and calibration on validation data could be explored to adjust the precision–recall balance.

First, while Pinder reduces redundancy via interface-based clustering, distribution shift across species, oligomerization states, and experimental conditions remains a challenge; explicit domain adaptation or contrastive debiasing may improve generalization. Second, our structural pathway assumes availability of reasonably accurate monomer structures. Although modern predictors provide broad coverage, uncertainty in flexible/disordered regions can degrade geometric signals; uncertainty-aware masking (e.g., pLDDT- or B-factor–guided) is a natural extension. Third, we froze ESM-2 for efficiency; parameter-efficient fine-tuning (e.g., LoRA [38]) is a possible avenue to test whether recall can be improved.

PeTriPPI2 illustrates the reuse of a structure-aware encoder trained on monomers for a pairwise prediction task. The code and model checkpoint availability information will be added here when the public release is ready.

PeTriPPI2 achieves competitive performance on Pinder while exhibiting a different precision–recall balance from SpatialPPIv2. The results motivate further evaluation of pretrained structural encoders across protein tasks. Future work will focus on (i) parameter-efficient tuning of the sequence pathway, (ii) uncertainty-aware geometric masking, and (iii) multi-task training that jointly learns *interaction* and *interface* signals.

## 5 Acknowledgments

Funding: Our work is publicly funded through ANR (the French National Research Agency) under the program named DeepPep ANR-23-CE20-0020-01.

This project was provided with computing HPC and storage resources by GENCI at IDRIS thanks to the grant 2025-AD010714598R2 on the supercomputer Jean Zay’s V100 and A100 partitions.

## 6 Data and code availability

- This work is based on the Pinder database ([32]) available at https://github.com/pinder-org/pinder
- PeTriPPI2 graph model: https://github.com/Baldwin-disso/PeTriBox.

## Notes

### Competing Interest Statement

The authors have declared no competing interest.

